# The Perceptual Neural Trace of Memorable Unseen Scenes

**DOI:** 10.1101/414052

**Authors:** Yalda Mohsenzadeh, Caitlin Mullin, Aude Oliva, Dimitrios Pantazis

## Abstract

Some scenes are more memorable than others: they cement in minds with consistencies across observers and time scales. While memory mechanisms are traditionally associated with the end stages of perception, recent behavioral studies suggest that the features driving these memorability effects are extracted early on, and in an automatic fashion. This raises the question: is the neural signal of memorability detectable during early perceptual encoding phases of visual processing? Using the high temporal resolution of magnetoencephalography (MEG), during a rapid serial visual presentation (RSVP) task, we traced the neural temporal signature of memorability across the brain. We found an early and prolonged memorability related signal recruiting a network of regions in both dorsal and ventral streams, detected outside of the constraints of subjective awareness. This enhanced encoding could be the key to successful storage and recognition.

## INTRODUCTION

Every day, we are bombarded by a mass of images in newspapers, billboards, and on social media, among others. While most of these visual representations are ignored or forgotten, a select few will be remembered. Recent studies have shown that these highly memorable images are consistent across observers and time scales demonstrating that memorability is a stimulus driven effect^1-5^. This is punctuated by the fact that observers have poor insight into what makes an image memorable. For instance, features such as interestingness, attractiveness and subjective memorability judgments (what the observer thinks they will remember) do not explain the phenomenon^2,6^.

Investigations into the neural basis of memorability using fMRI have revealed greater contributions in brain regions associated with high-level perception along ventral visual stream, rather than prefrontal regions associated with episodic memory^7-8^. These greater perceptual correlates indicate a potential processing advantage of memorable images, suggestive of a stronger perceptual representation. In a related vein, using an RSVP paradigm mixing images with different levels of memorability, Broers et al. (2017) found that memorable images were recognized significantly better than non-memorable images with extremely brief display durations^9^ (13ms), suggesting that features underlying image memorability may be accessible early on in the perceptual process. The early response of the memorability effect is also supported by work tracking pupillary response and blink rates for memorable and non-memorable images, concluding that memorable images have both an immediate^10^ and long-lasting^5^ effect on recognition performance.

Taken together, this work suggest that memorable images are encoded more fluently, and this perceptual processing advantage correlates with better long-term storage. Here, we trace the temporal dynamics of memorable images in order to reveal the time course of neural events that influence future memory behavior. We employed high temporal resolution of magnetoencephalography (MEG) during a rapid serial visual presentation (RSVP) task to isolate the perceptual signature of memorability across the brain.

In order to focus our investigation on purely perceptual aspects of memorability, we isolated the neural signal of memorable images from the influence of higher cognitive processes such as the top-down influence of memory. This approach requires the consideration of two basic principles: First, we acknowledge that image masking procedures, such as those found in traditional RSVP tasks, inhibit neural representations of non-target images from reaching awareness^11-16^. Second, we assume that memorability scores from the LaMem dataset^17^ (normative memory scores collected from thousands of observers) can function as a proxy for individual memory in the current study (i.e. an image with a high memorability score would be very likely to be remembered by an observer in our study had the information not been interrupted through masking, whereas an image with a low score would not)^2-5^.

Results revealed an early and prolonged memorability related signal recruiting a network of regions in both dorsal and ventral streams, detected outside of the constraints of subjective awareness. The enhanced perceptual encoding shown here could be the key to improving storage and recognition.

## RESULTS

During an ultra-rapid serial visual presentation (RSVP) paradigm^18^, observers performed a two-alternative forced-choice face detection task (Figure 1), while MEG data were collected. In each RSVP sequence of 11 images (34ms per stimulus), 15 participants were instructed to determine whether the middle image, or target, was a face or non-face (50% chance), a task they could perform successfully (d’ =3.72, two-sided signed-rank test; p<10^-4^). Importantly, in the face-absent trials the middle image was replaced by a scene image (the task-irrelevant target) sampled randomly from 30 images, half with a high-memorability score of 0.88±0.06 (mean±std) and half with a low-memorability score of 0.59±0.07 (mean±std). The remaining images (distractors) in the sequence were sampled from mid-level memorability scores of 0.74±0.01 (mean±std). Task-irrelevant target and distractor stimuli came from the LaMem dataset, with pre-acquired memorability scores^17^.

**Figure 1.**
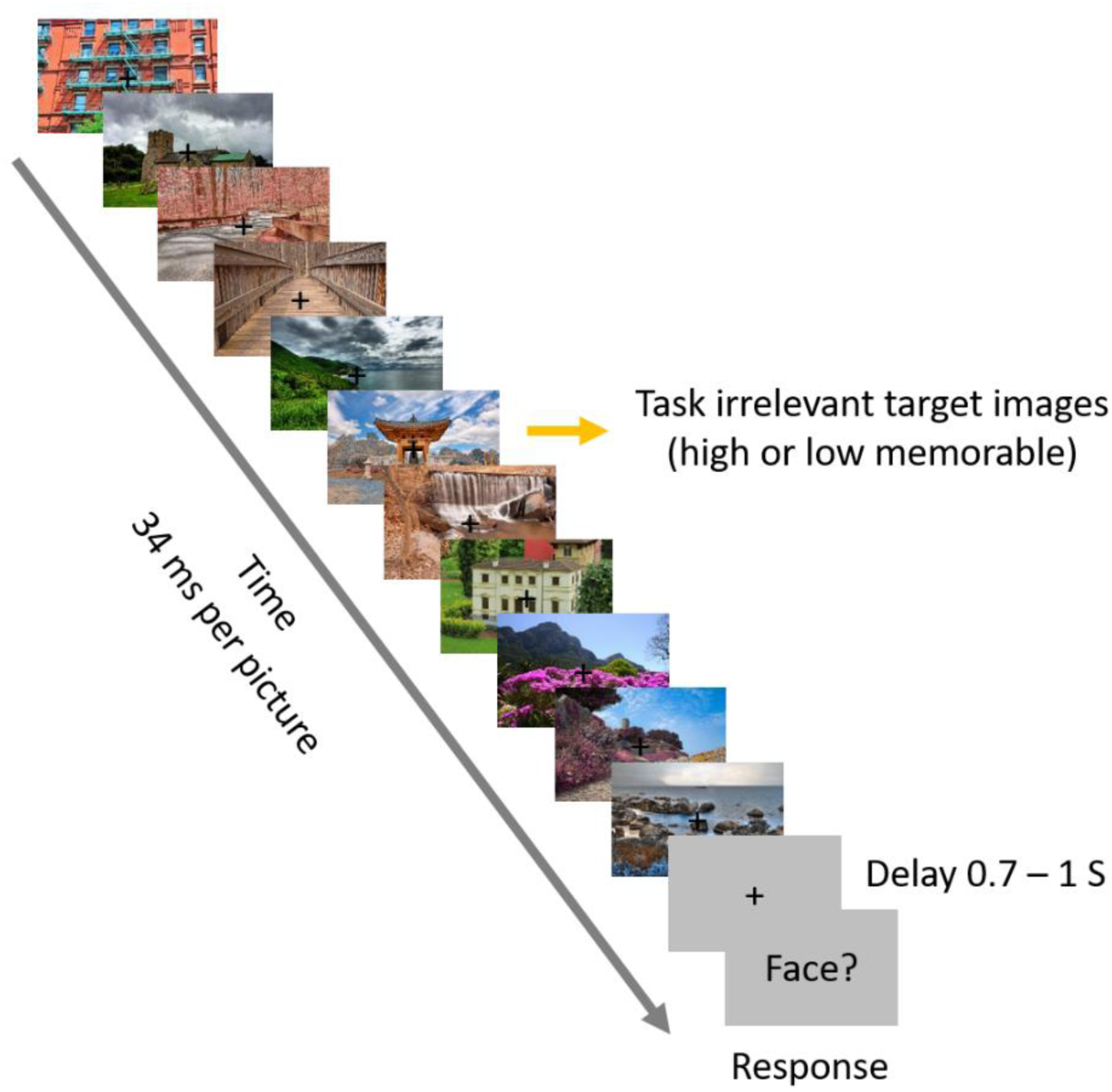
Paradigm design. RSVP paradigm, known to greatly reduce awareness to images, and examples of high and low memorable task-irrelevant scene targets. Each RSVP trial includes presentation of 11 images with the speed of 34 ms per picture (without inter-stimulus interval). In half of the trials a face image was embedded in the middle of the sequence and participants were asked to detect the face trials (a two-alternative forced choice task). In the non-face trials, the middle image was drawn randomly from a set of 30 scene images, half high memorable and half low memorable (task-irrelevant targets). The presented images in this figure are not examples of the stimulus set due to copyright. Depicted images are in publicly available at flicker.com under a Creative Commons (CC) license.

Following the MEG experiment, observers performed an unanticipated old-new memory test, with all targets and novel images shown one at a time. Participants were presented with the RSVP target images mixed randomly with 60 novel images matched on high and low level features, and instructed to indicate whether they had seen the image at any point during the RSVP task. Results show that while the goal-directed target stimuli (faces) were detected and remembered well above chance (d’ = 0.59, two-sided signed-rank test; p < 0.01), memory for task-irrelevant target scenes was at chance level (d’ = - 0.13, two-sided signed-rank test; p = 0.1), corroborating an absence of explicit memory trace. The unanticipated memory tests confirmed that despite 30 repetitions (see methods) of each task-irrelevant target scene, participants failed to recall having seen these images. This provides the ideal circumstances to evaluate the purely perceptual basis of memorability, thus all subsequently described analyses focused only on perceptual dynamics of the task-irrelevant scene stimuli and the consciously perceived face-target trials were disregarded from further analyses.

### Temporal trace of memorable images

After extracting the MEG time series from -100 to 500ms relative to task-irrelevant scene target onset, we performed multivariate pattern analysis in a time-resolved manner. For each time point (1ms step), we measured the performance of a SVM classifier to discriminate between pairs of scene images using leave-one-out cross-validation resulting in a 30 x 30 decoding matrix, also known as representational dissimilarity matrix (RDM), at each time point (Figure 2A). We then used the representational similarity analysis (RSA) framework^19-22^ to characterize the representational geometry of memorability effect in MEG data. In this framework, hypothesized model RDMs can be compared against time resolved RDMs created from MEG data by computing their correlations (Figure 2A). Here we considered two hypotheses, a linearly separable representation between our two conditions, such as a categorical clustering geometry (see the model RDM and its 2D multidimensional scaling (MDS) visualization in Figure 2B), and a nonlinear entropy based representation where one condition is dispersed while the other is tightly clustered (see the model RDM and its 2D MDS visualization in Figure 2C). The comparison of these two candidate model RDMs with the time resolved MEG RDMs yielded the correlation time series presented in Figure 2B and C. As depicted, while no significant correlations were found between the MEG RDMs and the linearly separable model in Figure 2B, the model assuming a more entropy based geometrical representation for high memorable images explained MEG RDM patterns with significant correlations starting at ∽150 ms after target image onset.

**Figure 2.**
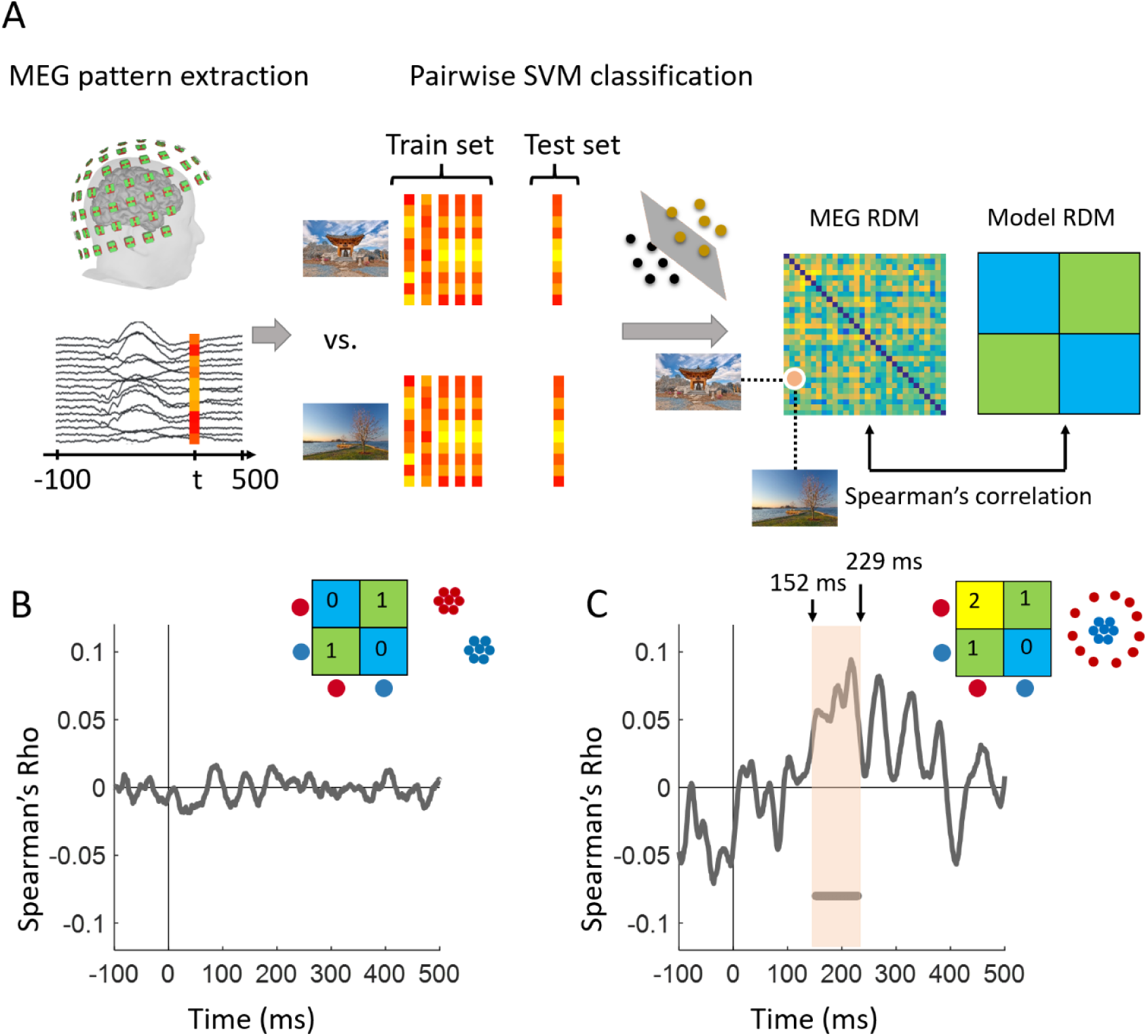
Multivariate pattern analysis and geometrical representation of memorability across time. (A) Using MEG pattern vectors at each time point t, a support vector machine (SVM) classifier was trained to discriminate pairs of target scene images. The performance of the SVM classifier in pairwise decoding of target images populated a 30 × 30 decoding matrix at each time point *t*. This process resulted in time resolved representational dissimilarity matrices for MEG data which can then be compared with candidate model RDMs by computing their Spearman’s rho correlations. (BC) Two possible representational geometries of memorability and their comparison with MEG data. The RDM and MDS plot in panel B show a categorical representation in which high and low memorable images are linearly separable. The RDM and MDS plot in panel C illustrate a representational geometry where high memorable images are more dispersed than low memorable ones. The gray curves in (B) and (C) depict the time course of MEG and model RDM correlations. The line below the curve in panel C indicates significant time points when the correlation is above zero. Statistical tests used a cluster-size permutation procedure with cluster defining threshold *P*<0.05, and corrected significance level *P*<0.05 (n=15).

The lack of categorical separability in our representational geometry implies that the classical between-categorical decoding analysis is not well suited to describe the distinction between these two conditions. Instead, we averaged the decoding values (dissimilarities) within high and low memorable scene pairs separately. Figure 3 shows the two curves for high and low memorable scenes across time in red and blue, respectively. Decoding accuracy was near identical for high and low memorable scenes until 149ms, at which point the two curves diverged significantly, revealing the onset of a memorability-specific signal, which lasted until 228ms consistent with the results in Figure 2C.

**Figure 3.**
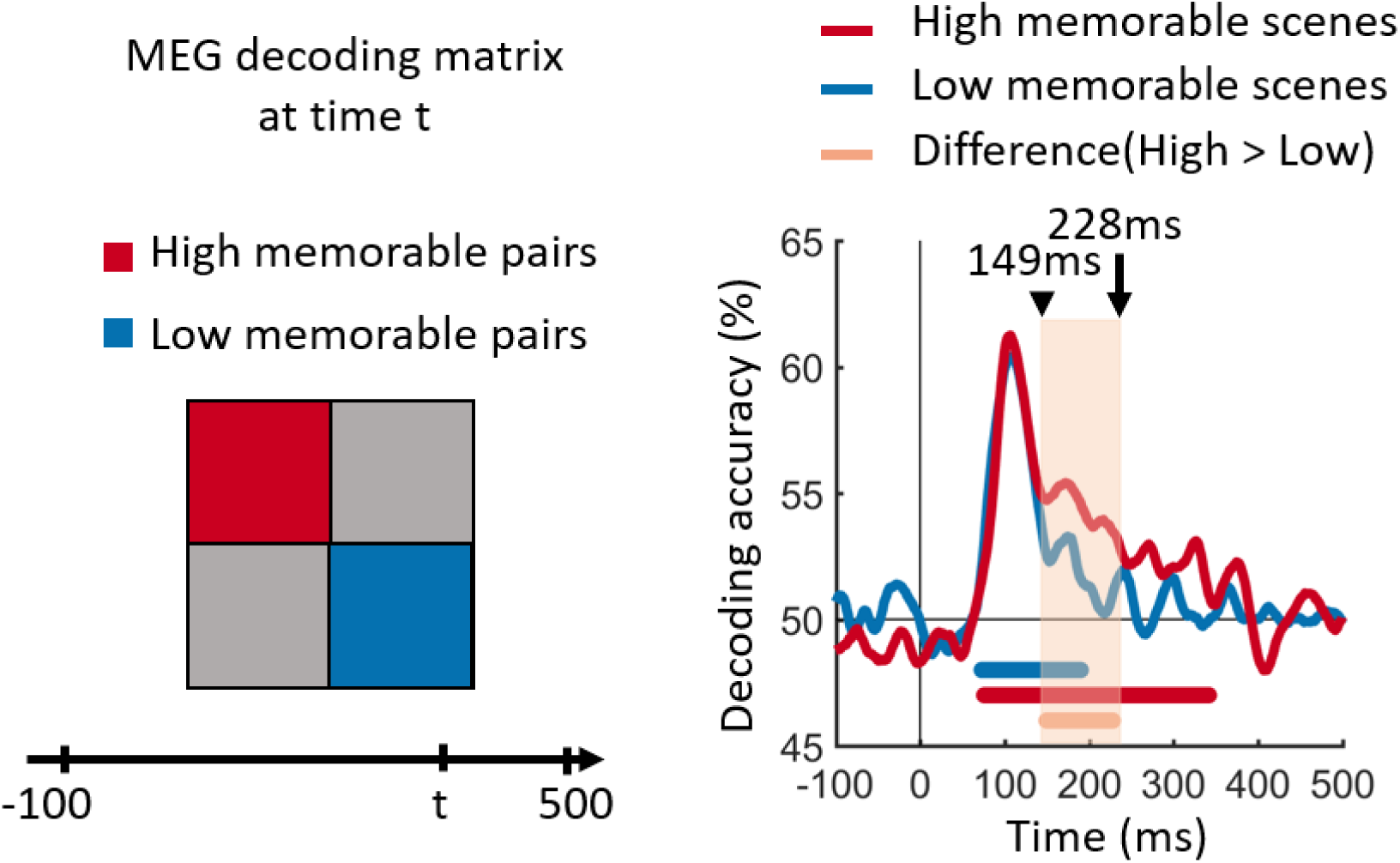
Time course of image decoding for high versus low memorable images. The pairwise decoding values were averaged within high and low memorable images separately. The color coded red and blue lines at the bottom of curves show significant time points where the decoding is above the chance level of 50%. The orange line indicates significant time points for the difference between high and low memorability. Statistical tests used a cluster-size permutation procedure with cluster defining threshold *P*<0.05, and corrected significance level *P*<0.05 (n=15).

Together, our analysis demonstrates that the two categories of high and low memorable images are not linearly separable, but that high memorable images show a more distributed geometrical representation than low memorable images. This suggests that memorable stimuli are associated with higher differentiability and unique information, as illustrated by higher averaged decodings within high memorable scene pairs than low memorable scene pairs (Figure 3).

### Temporal generalization reveals an evolutionary dynamics for memorable images

The significant persistence of memorable images from 149-228ms (cluster defining threshold *p*<0.05; corrected significance level *p*<0.05) suggests this class of stimuli benefited from prolonged temporal processing. This extended processing could manifest as either a stable representation sustained over time - i.e. as a form of neural maintenance, or a series of distinct representations dynamically evolving over time. To investigate, we applied a temporal generalization approach^23^ which uses the trained SVM classifier on MEG data at a given time point t (training time) to test on data at all other time points t’ (testing time). Intuitively, if neural representations sustained across time, the classifier should generalize well across other time points. The resulting matrices (Figure 4AB), in which each row corresponds to the time (in ms) at which the classifier was trained and each column to the time at which it was tested, reveal that both conditions show a diagonally extended sequence of activation patterns starting at ∽ 70ms. This shape of significant time points suggests that the representations of both conditions dynamically evolved over time. Importantly, the greater diagonal reach of the high memorable condition suggests further processing during this evolutionary chain.

**Figure 4.**
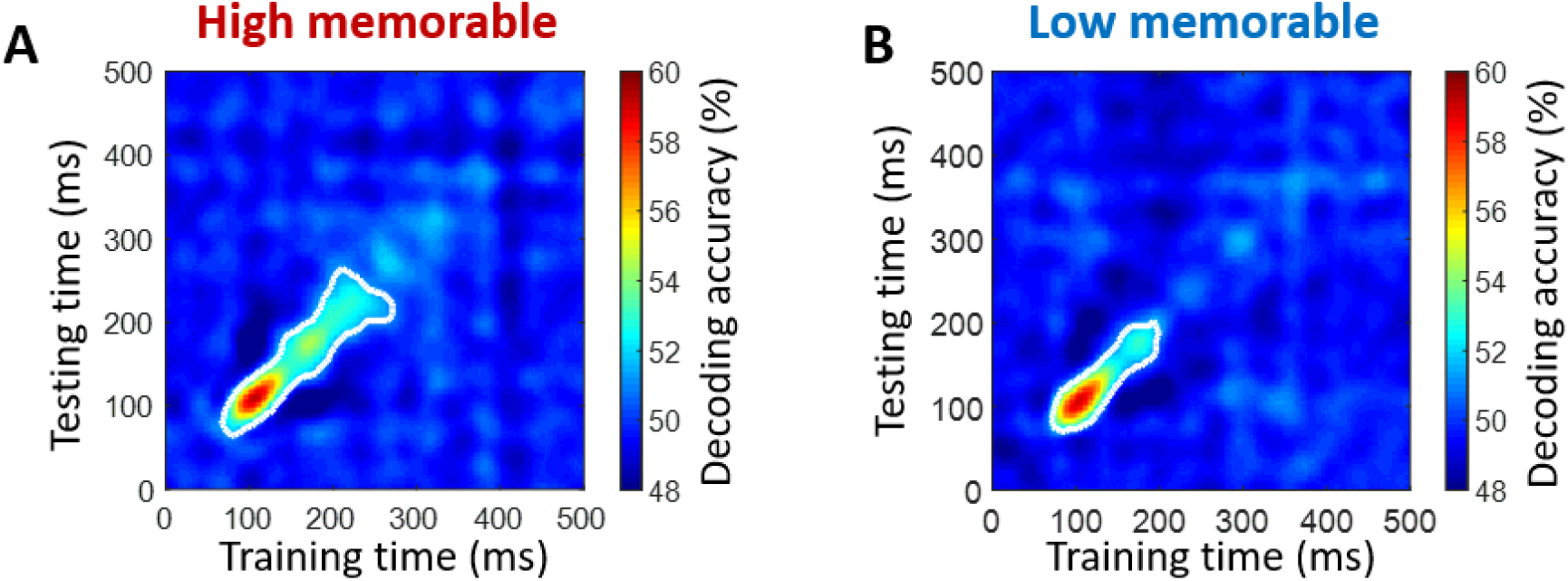
Temporal generalization. (AB) Generalization of image decoding across time for high and low memorable images. The trained SVM classifier on MEG data at a given time point t (training time) was tested on data at all other time points t’ (testing time). The resulting decoding matrices were averaged within high or low memorable scene images and over all subjects. White contour indicates significant decoding values. Statistical tests used a cluster-size permutation procedure using cluster defining threshold *P*<0.05, and corrected significance level *P*<0.05 (n=15).

### Cortical source of image memorability

How do the changing patterns of brain activity map onto brain regions? Using MEG source localization, we investigated where the memorability-specific effects manifested in the visual stream. We broadly targeted three anatomically defined regions-of-interest (ROI) known to be involved in building visual image representation. Based on Freesurfer automatic segmentation^24^, we selected the pericalcarine area for early visual processing, the inferior temporal area for the ventral stream processing and a parietal area for the dorsal stream processing. Within these ROIs we performed the same pairwise decoding analysis as for the sensor data but now using cortical source time series within these regions. Decoding results suggest that memorability recruited distinct brain regions evolving over time: as expected, no significant differences were seen in the pericalcarine (Figure 5AD), while significant differences between high-and low-memorable images were localized in the left parietal area starting at 153ms (Figure 5B) and then later in the right inferior-temporal around 225ms (Figure 5F).

**Figure 5.**
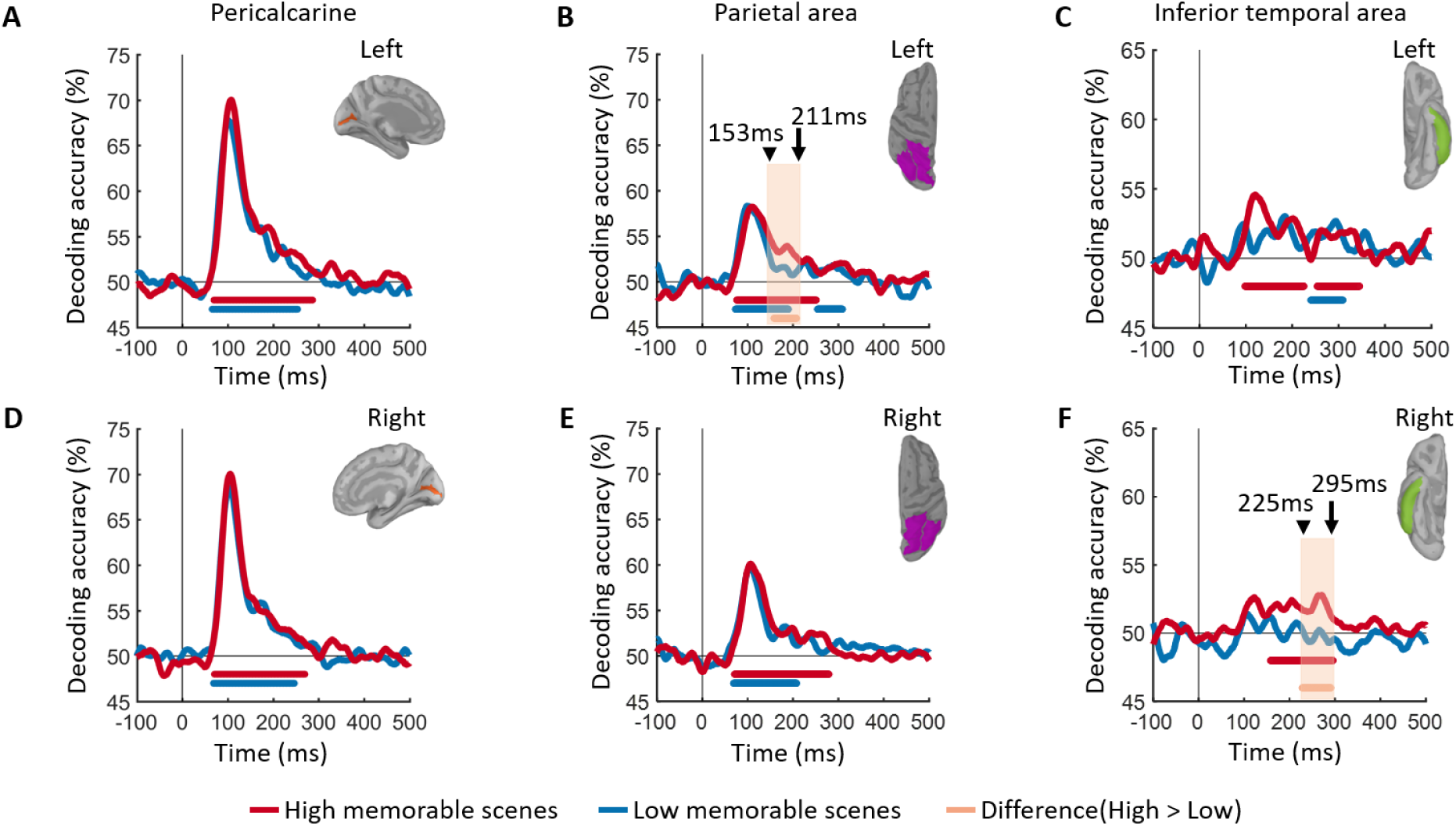
Spatial localization. (A-F) Time course of image decoding in cortical sources. RDM matrices were extracted at each time point t using MEG source localization in pericalcarine, parietal area and inferior temporal, separately for left and right. The color coded red and blue lines at the bottom of curves show the significant time points where the decoding is above chance level of 50%. The orange line indicates significant time points for the difference between high and low memorability. All significant statistical tests are with permutation tests using cluster defining threshold *P*<0.05, and corrected significance level *P*<0.05 (n=15).

## DISCUSSION

In the current study, we examined the temporal processing signature of visual information that is likely to be remembered compared to the one likely to be forgotten. We tested this effect using high-temporal resolution MEG during an RSVP task to suppress the effects of top-down influences of memory. Our results revealed the dynamic neural blueprint of the perceptual memorability effect, with highly memorable images showing significantly better decoding accuracies between ∽150-230 msec. Despite the extremely rapid viewing conditions, this signal persisted and evolved over multiple brain regions and timescales associated with high-level visual processing (e.g. semantic category or identity information).

A variety of neuroimaging and recording techniques have demonstrated that the cortical timescale of visual perception begins with low level features in early visual cortex at ∽40-100 msec^24-29^ and reaches the highest stages of processing in the inferior temporal cortex within 200 ms after stimulus onset^21-22,30-37^. Despite memorability being a perceptual phenomenon, previous work has revealed that low-level image features, as well as non-semantic object statistics, do not correlate strongly with memorability scores^2^, thus these features were equalized between conditions here, resulting in the overlapping curves within the first 150 msec after image onset (Figure 3) and the lack of any memorability effects in the pericalcarine ROI (Figure 5AD).

Event-related potentials (ERPs) have linked high-level properties of the visual stimulus, such as its identity or category, with a timescale roughly 150 msec after stimulus onset^30-31^. More recently, MEG work has demonstrated a similar pattern of results^21-22,38^. While the current image set also controlled for high-level semantics between conditions (for every high memorable stimulus, there was a matching low memorable stimulus of the same semantic category), the memorability effect persisted during this period such that the decoding rate for memorable images did not drop as drastically, but instead persisted for an additional ∽100 msec (see Figure 3). This suggests that the timescale of the memorability effect (∽150-230 msec) is reflective of a processing advantage for high level perceptual features.

How does this persistent high-level signal reflect the processing advantages of memorable images? To address this question, we examined the temporal generalization of the MEG data to reveal how the sequence of the memorability signals manifests over time. While the task-irrelevance and rapid masking in the current design inhibited the stimulus representations from reaching awareness, recent temporal generalization work suggests that unseen stimuli are still actively maintained in neuronal activity over time, with an early signal representing the evolution of perception across the visual hierarchy (diagonal pattern) and a later signal generalizing over time (square pattern) as a function of the subjective visibility of the stimulus^39^.

Given that high memorable images are quickly perceived and understood^9^, we might have expected the memorable representations to stabilize more rapidly in the brain (as evidenced by a square generalization matrix), potentially reflecting a more durable maintenance of information across time. However, in both the high and low memorable conditions the generalization matrices were dominated by a diagonal pattern, commonly associated with a long sequence of neural responses reflecting the hierarchical processing stages of perception^23^. Importantly, the high memorable representations demonstrated a significant diagonal extension over the low memorable condition (Figure 4), suggesting greater evolution of perception during this processing chain. Thus, the prolonged diagonal shape suggests that rather than manifest into memories under the current rapid viewing conditions, the representations of these memorable images were strong enough to take one step further down the perceptual processing pathway, perhaps readying themselves for a reliable transition into long term storage.

The evolution of signal observed in the diagonal processing chain triggers the question of which brain regions are responsible for this perceptual advantage? We performed an ROI based cortical source localization to evaluate the contributions of several regions previously associated with perceptual processing. Results revealed that memorability was localized to several high-level brain regions evolving over time; with the left parietal region recruited first, followed by the right IT cortex and, as predicted, no-significant difference observer in the pericalcarine. While hemispheric laterality effects may be a reflection of low signal-to-noise ratio, our results implicate high-level visual cortex in both the dorsal and ventral streams and not early visual areas to memorability effects.

The significantly better decoding rate of high-memorable images in these brain regions indicates that their cortical representations reflect a more concrete embodiment of the stimulus compared to those of low-memorability. However, this strong representation is still evolving over time and space as it moves from one high-level perceptual region to another. Both the inferior-temporal region and areas encompassed by the parietal cortex have been previously associated with the perception of shape, objects, faces and scenes and other high-level visual features^40-46^. Thus, when these regions are recruited during natural viewing, memorable images seem to carry a hidden advantage in the form of a kind of processing fluency^47^.

As the medium of knowledge communication continues to evolve, visual literacy (the skill to interpret, negotiate, and make meaning from information presented in the form of an image) has become increasingly important. The ability of some images to be quickly understood and stick in our minds provides a powerful tool in the study of neural processing advantages leading to superior visual understanding. The high temporal-resolution results reported here provide new insights into the enduring strength of perceptual representations, pinpointing a high-level perceptual property that is quickly encoded.

While high-memorable images are more likely to be subsequently remembered, the masking effects of the RSVP paradigm halted the processing of perceptual information in our study and prevents us from making any claims about the transition from strong perceptual representation to memory. Future work should examine the full timeline of memorability in the brain; from encoding, to storage to recognition. Given the low spatial resolution of MEG, future work should also focus on linking higher spatial resolution brain data to the current timescale. The ability to reliably trace the memorability signal over space and time has many practical advantages such as the early detection of perception or memory impairments in clinical populations (weaker or slower representations^48^ and the design and establishment of more memorable educational tools for improved implicit visual literacy.

### METHODS

### Participants

Fifteen healthy right-handed human subjects (12 female; age mean ± s.d. 23.8 ± 5.7 years) with normal or corrected to normal vision participated in this experiment after signing an informed written consent form. They all received payment as a compensation for their participation. The study was approved by the Institutional Review Board of the Massachusetts Institute of Technology and conducted in agreement with the principles of the Declaration of Helsinki.

### Experimental Design and Stimulus set

#### Stimulus set

The stimulus set comprised 60 target images (30 faces and 30 scenes) and 150 distractor images of scenes. The scene images (task irrelevant targets and distractors) were selected from a large memorability image dataset called LaMem^17^. The 30 scenes comprised of 15 high memorable and 15 low memorable images controlled for low level features (color, luminance, brightness, and spatial frequency) using the natural image statistical toolbox^49^. The face target images were selected from the 10K USA Adult Faces^3^.

#### RSVP paradigm

Participants viewed RSVP sequences of 11 images each presented for 34ms without inter stimulus interval in separate trials (Figure 1). The middle image and the 10 distractor images, respectively, were randomly sampled from the set of 60 target images and the set of 150 distractor images. The image sequence was presented at the center of the screen on a gray background with 2.9° of visual angle. Each trial started with a 500ms baseline time followed by the RSVP sequence and then a blank screen which was presented for 700 – 1000ms with uniform distribution. The blank screen aimed to delay response and thus prevent motor artifacts on the data. At the end of trial, the subjects were prompted with a question to report whether they have seen a face image in the sequence or not and they responded with their right thumb using a MEG-compatible response button box. The experiment included 30 trials for each of the target images and trials were randomly ordered and presented in 12 blocks with 150 trials. In order to prevent eye movement artifacts, participants were instructed to fixate on a black cross at the screen center and only blink when pressing the button to respond. The subjects did not see the target images or distractors before the experiment.

#### Subsequent memory test

To determine if the target images (middle images in each sequence) were encoded in memory or not, after the MEG experiment we asked the subjects to perform an unanticipated memory test. They were presented with 120 images, the 60 RSVP target images randomly mixed with 60 novel images (30 faces and 30 scenes matched in terms of low level features and semantics with the target images), and were asked to report if they have seen the images during the experiment with 4 levels where 1 being a confidently novel image and 4 being a confidently seen image.

### MEG acquisition and preprocessing

MEG data was collected using a 306-channel Elekta Triux system with the sampling rate of 1000 Hz and a band-pass filter with cut-off frequencies of 0.03 and 330 Hz. We measured the head position prior to the MEG recording using 5 head position indicator coils attached to the subjects’ head. The head position was also recorded continuously during the experiment.

Maxfilter software was applied on the acquired MEG data for head movements compensation and denoising using spatiotemporal filters^50-51^. Then Brainstorm software^52^ was used to extract trials from 100ms before to 500ms after target image onset and preprocess the data. We removed the baseline mean of each sensor and data was smoothed by a low-pass filter of 20Hz. Trials with amplitude greater than 6000 fT (or fT/cm) were marked as bad trials. Eye blink/movement artifacts were detected using frontal sensor MEG data and then principal component analysis was applied to remove these artifacts from the MEG data.

### MEG multivariate pattern analysis

#### Sensor space

We analyzed MEG data using multivariate pattern analysis. To decode information of the task irrelevant target stimuli, a linear support vector machine (SVM, libsvm implementation^53^) was used as a classifier. In order to reduce computational load, the MEG trials of each condition were sub-averaged in groups of 5 with random assignment, resulting in N = 6 trials per condition. At each time point t of each trial, the MEG data was arranged in a vector of 306 elements. Then, for each pair of high or low memorability scene images (middle scenes in the RSVP sequence) and at each time point, the accuracy of SVM classifier was calculated using a leave-one-out procedure. The procedure of sub-averaging and then cross-validation was repeated for 100 times. The classifier accuracies were averaged over the repetitions separately for pairs of high or low memorability scene images.

#### Source space

To localize representational information on regions of interest (ROIs), we mapped MEG signals on cortical sources (based on Freesurfer automatic segmentation^24^ using default anatomy^54^) and performed multivariate pattern analysis on each ROI. We computed the forward model using an overlapping spheres model^55^ and then using a dynamic statistical parametric mapping approach (dSPM) MEG signals were mapped on the cortex^56^. Time series from cortical sources within three regions of interest, namely, pericalcarine, inferior temporal and parietal area (concatenating inferior parietal and superior parietal) were derived^57^. In each cortical region of interest, pattern vectors were created by concatenating ROI-specific source activation values, and then a similar multivariate pattern analysis was applied to the patterns of each ROI.

### Temporal generalization with multivariate pattern analysis

To compare the stability of neural dynamics of high and low memorable images, we studied the temporal generalization of their representations^21-23, 60^ by extending the SVM classification procedure. The SVM classifier trained at a given time point t was tested on data at all other time points. The classifier performance in discriminating signals can be generalized to time points with shared representations. This temporal generalization analysis was performed on every pair of images and for each subject. Then averaging within high (or low) memorable images and across subjects resulted in a 2D matrix where the x-axis corresponded to training time and y-axis to testing time.

### Statistical inference

We used nonparametric statistical tests which do not assume any distributions on the data^61-62^. Our statistical inference on decoding time series and temporal generalization matrices were performed by permutation-based cluster-size inference (1000 permutations, 0.05 cluster definition threshold and 0.05 cluster threshold) with null hypothesis of 50% chance level. For difference of decoding time series we used 0 as chance level. We performed bootstrap tests to assess statistics for peak latency of time series. Specifically, we bootstrapped subject-specific time series for 1000 times, each time we averaged the time series and found its peak latency, and finally using the empirical distribution of peak latencies we assessed the 95% confidence intervals.

## AUTHOR CONTRIBUTIONS

Y.M., A.O. and D.P. designed the experiment. Y.M. conducted the experiment and analyzed the data. Y.M., C.M., A.O. and D.P. interpreted the results and wrote the manuscript.

## ACKNOWLEDGMENTS

This work was funded by NSF award 1532591 in Neural and Cognitive Systems (to D.P and A.O) and the Vannevar Bush Faculty Fellowship program by the ONR to A.O. (N00014-16-1-3116). The study was conducted at the Athinoula A. Martinos Imaging Center at the McGovern Institute for Brain Research, Massachusetts Institute of Technology.

